# *Drosophila* Arc1 is required for metabolic resilience and developmental timing during dietary challenges

**DOI:** 10.64898/2026.04.14.718561

**Authors:** Wei Zhang, Lauren Schmitt, Kent Riemondy, Tânia Reis

## Abstract

Energy homeostasis at the organismal level requires balancing energy storage and mobilization to provide sufficient fuel for energy-intensive processes like development without depleting or accumulating excess stores. Fluctuations in the nutritional content of the diet present a challenge to the pathways that maintain energy balance. We previously identified the *Drosophila melanogaster* counterpart of human ARC (activity-regulated cytoskeleton-associated protein) as a brain-expressed protein that regulates energy storage in the major fat storage tissue of the fly, the fat body. Here we show that Arc1 expression in the brain responds to changes in diet and insulin-like peptide levels. Mutating Arc1 perturbs the ability of larvae to maintain normal body fat and rates of development upon dietary changes: mutants develop slower or faster than wild-type on nutrient-poor or nutrient-rich diets, respectively. Excess fat storage in *Arc1* mutants becomes an advantage upon starvation, prolonging survival relative to the wild type. In addition to metabolic and neuronal genes, transcriptomic analysis revealed changes in key developmental drivers of development, in both diet-dependent and - independent manners. This study supports a model in which nutrient regulation of Arc1 via insulin-like peptide signaling couples dietary changes to changes in metabolism –– to maintain energy homeostasis –– and production of hormone signals, to support timely development. In this role, Arc1 is a central player in a buffering mechanism that coordinates nutrient availability, organismal metabolism, and developmental rate.

## Introduction

Homeostasis is a defining feature of life. Energy homeostasis involves a balance between usage, storage, and liberation of energy. Energetic requirements –– and therefore challenges to energy homeostasis –– change during development and lifespan (Gillette et al. 2021). As in many species, in *Drosophila melanogaster* different developmental stages exhibit distinct energetic demands (Flecka et al. 2025). Embryonic development is a closed system fueled by maternally inherited resources. The larval stage is marked by rapid, energy-intensive cellular growth and proliferation fueled by anaerobic glycolysis (Drummond-Barbosa and Tennessen 2020; Mattila and Hietakangas 2017). Pupal development is another closed system with no feeding inputs, relying solely on internal stores acquired during the earlier larval feeding period. As with most animals, when flies reach adulthood most energy-intensive growth ceases, and nutritional requirements are coupled instead to behaviors that ensure survival and reproduction (Flecka et al. 2025). Accordingly, energy intake (i.e., diet quantity and quality) must be carefully controlled across development to provide sufficient fuel to sustain growth and survival while maintaining appropriate energy stores.

Progression through development is controlled by multiple endocrine signals. During larval development, the steroid hormone ecdysone (Ec) is a critical player coordinating development and growth with available resources (reviewed in (Yamanaka et al. 2013a)). Ec biosynthesis requires dietary cholesterol to be metabolized in the prothoracic gland. Changes in diet modulate Ec synthesis and release (reviewed in (Pan et al. 2021). Ec activity is regulated at different levels by other hormones. The secreted peptide Prothoracicotropic hormone (PTTH) is a major regulator of Ec (Pan et al. 2021), stimulating Ec synthesis by binding to a receptor tyrosine kinase and initiating downstream ERK signaling (Rewitz et al. 2009). Ec synthesis is repressed by juvenile hormone (JH), levels of which spike during larval development to inhibit precocious metamorphosis (Liu et al. 2018). It is the antagonistic coordination of Ec and JH hormones that result in the three molts associated with the transitions between larval instars, and ultimately metamorphosis (reviewed in (Santos et al. 2019). Insulin-like peptides (ILPs) and associated receptors respond to nutrients and integrate coordination of growth with development (Nassel and Vanden Broeck 2016). Accordingly, variation in diet can alter the pace of development; since nutritional availability can change unpredictably, feedback loops involving a number of pathways and check points ensure resilience in this process. These loops ensure that, within limits, animals progress through the different instars despite oscillations to nutritional availability while continuing to store the extra energy required for metamorphosis. More dramatic changes to diet ––restriction or excess of all and/or specific nutrients –– compromises developmental timing, energy storage levels, growth rate, fertility and/or viability (Martelli et al. 2024; Reis 2016; Sorge et al. 2025). Of note, custom diets designed to match the metabolism of mutant animals partially ameliorate the developmental delays, energy storage defects, and poor survival associated with mutating a conserved regulator of organismal metabolism (Gillette et al. 2020). Despite numerous studies, the mechanisms that buffer the speed of development in response to dietary changes are still not fully understood.

Here we report roles for Arc1, a neuronal regulator of synaptic plasticity, in maintaining energy balance during larval development and buffering against the effects of imbalanced diets. Arc1 was identified by sequence similarity to human ARC, a neuronal protein that is transcriptionally upregulated by synaptic activity and is required for synaptic plasticity (Bramham et al. 2010). *Drosophila* Arc1 is also induced by synaptic activity (Guan et al. 2005; Mattaliano et al. 2007) and Arc1 mutants exhibit synaptic plasticity defects (Ashley et al. 2018). An early phenotype observed in *Arc1*-mutant flies was a behavioral defect in response to starvation (Mattaliano et al. 2007). Like almost all other animals, when starved, adult *Drosophila* exhibit a burst of locomotor activity, presumably as a food-seeking behavior, but *Arc1* mutants actually move less when starved and are resistant to starvation (Mattaliano et al. 2007). Arc1 expression in insulin-like-peptide-producing neurons of the brain is sufficient to restore a normal response to starvation (Mattaliano et al. 2007). This observation provided the first evidence of a connection between Arc1 and energy homeostasis.

We subsequently identified *Arc1* in an unbiased genetic screen for mutants that increase fat storage in *Drosophila* larvae (Reis et al. 2010). Independently, we performed an unbiased neuronal ablation screen for regions of the larval brain that control fat storage, and found that Arc1 transcript levels change when we manipulate activity in one region, called E347 (Mosher et al. 2015). These findings led us to overexpress or deplete Arc1 and show that excess Arc1 is sufficient to decrease fat storage, while Arc1 depletion drives fat accumulation and prevents the lean phenotype caused by E347 hyperactivation (Mosher et al. 2015). In no case was Arc1 manipulation associated with changes in feeding or locomotor activity (Mosher et al. 2015). These results pointed to a role for Arc1 in the brain in controlling whole-organism metabolism. In support of our hypothesis, we found changes in levels of key metabolites and metabolic enzyme in *Arc1* mutants (Mosher et al. 2015). *Arc1*-mutant adults were subsequently found to also have more stored fat (Swope et al. 2023) and a high-throughput RNAi-based screen in adult flies also independently found increased adiposity associated with depletion of Arc1 (Pospisilik et al. 2010). Also consistent with its metabolic roles, Arc1 expression depends on the status of the microbiota: germ-free *Arc1*-mutant larvae experience a developmental delay, and the yeast content of the diet influences the extent of the delay (Keith et al. 2021).

Here we investigated the role of Arc1 across dietary challenges during development. We find that Arc1 expression changes in a diet-specific manner, and found a range of dietary conditions that affect *Arc1* mutants in opposing ways, i.e., by accelerating or delaying development. *Arc1* mutants misregulate *Jhe* (juvenile hormone esterase), which encodes a critical enzyme required for JH degradation and development progression (Crone et al. 2007; Hopkins et al. 2019; Liu et al. 2008), independent of diet. We propose that, together with the diet-specific metabolic and neuronal changes, the Jhe defect leads to broader variations in developmental timing in *Arc1* mutants compared to wildtype animals. We thus propose that Arc1 is necessary for developmental resilience upon dietary challenges.

## RESULTS

### Insulin-like peptide signaling decreases Arc1 brain expression

Exogenously added insulin induces ARC expression in cultured human cells (Kremerskothen et al. 2002), and *Drosophila* Arc1-expressing cells overlap with insulin-producing cells (IPCs) (Keith et al. 2021; Mattaliano et al. 2007). Thus we previously proposed that insulin-like peptide (ILP) levels in insulin-producing cells (IPCs) control Arc1 expression (Mosher et al. 2015). To directly examine the relationship between ILP levels and Arc1 expression, we overexpressed Ilp2 in IPCs and used immunostaining to count the number of Arc1-positive cells in the brain. Ilp2 overexpression reduced the number cells that expressed Arc1 in the subesophageal zone (SEZ) and ventral nerve zone (VNZ) (Figure 1A). Notably, despite the overexpression, Ilp2 protein levels were not detectably changed in IPCs (Figure 1A), suggesting that the Ilp2 produced in those cells was efficiently secreted. By contrast, we previously observed Ilp2 accumulation in IPCs upon E347 neuronal silencing and attributed this increase to inefficient secretion (Mosher et al. 2015). Thus Arc1 expression in *Drosophila* larvae responds to ILP signaling, specifically to Ilp2.

**Figure 1.**
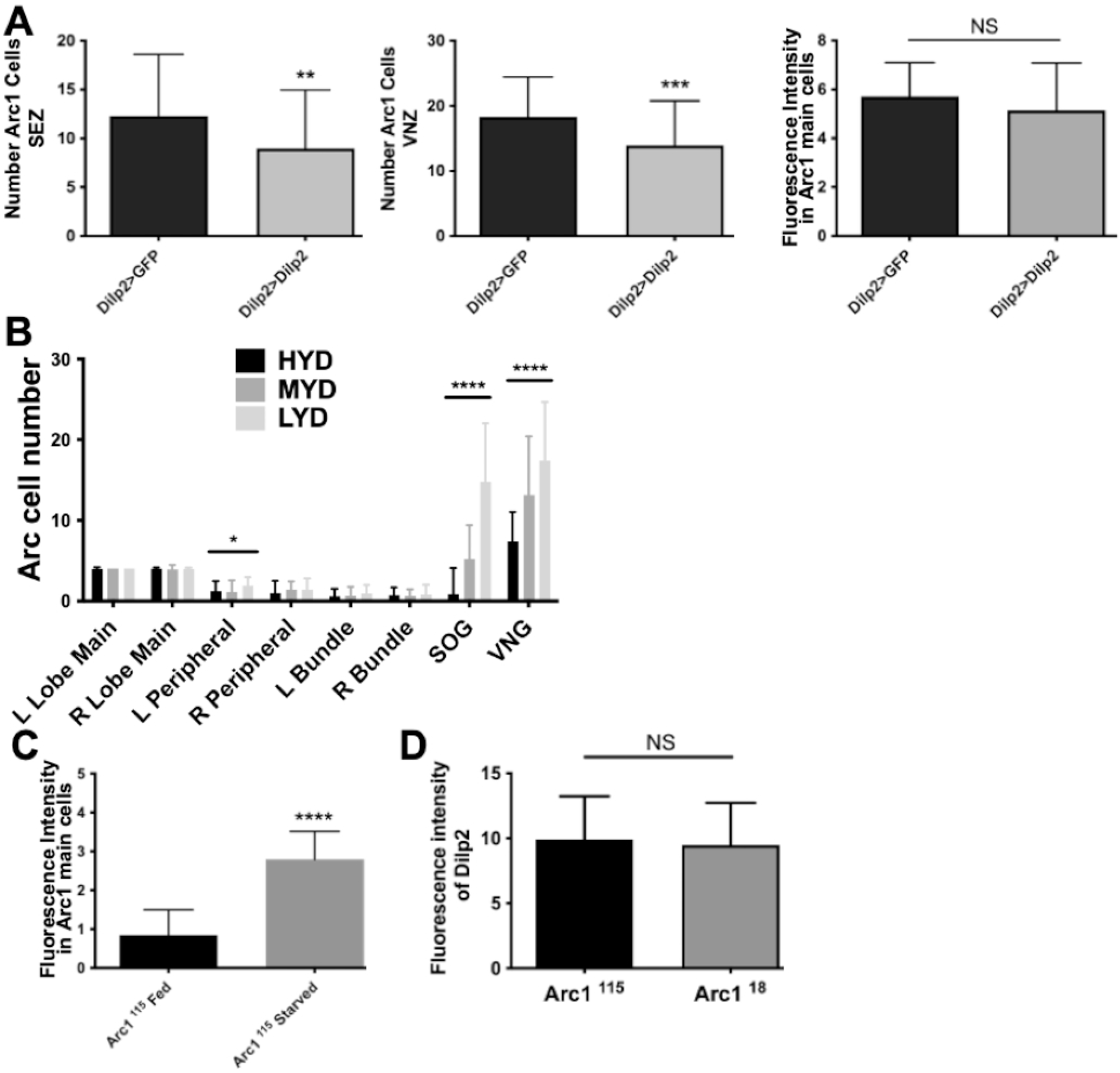
Arc1 expression in the brain responds to Ilp2 and diet. (A) Number of Arc1-positive cells in the indicated brain regions of larvae overexpressing GFP (as a control) or Ilp2. Larvae were fed a medium-yeast diet (MYD). n=50 brains in each condition from 3 independent biological replicates. P values calculated using two-tailed unpaired t test. (B) Number of Arc1-positive cells in the indicated brain regions of control (w^1118^) larvae fed high-yeast diet (HYD), MYD, or low-yeast diet (LYD). L, left lobe; R, right lobe; SEZ subesophageal zone; VNZ, ventral nerve zone. n=33-38 brains each condition from 3 independent biological replicates. Multiple t test comparison was used to calculate significance. (C) Levels of Arc1 protein in the Arc1-producing cells from larvae fed or starved for 12 hrs prior to dissection. Larvae were fed MYD until 84 hr after egg deposition, at which point they were either transferred onto agar-only medium (for starvation) or into new vials of food. n=19-31 brains from 3 independent biological replicates. P values calculated using two-tailed unpaired t test. ***, 0.0001<P< 0.001; **, 0.001<P<0.01; *, 0.01<P<0.05.

### Arc1 levels in the brain change in response to diet

ILP expression is tightly coupled to nutrition: Ilp2 responds to levels of yeast in the diet (Suzawa et al. 2025). Considering the effect of Ilp2 expression we observed on Arc1 levels, we asked if Arc1 expression might also respond to changes in levels of dietary yeast. We created a high-yeast diet (HYD) with twice the yeast content of the standard diet (medium-yeast diet, MYD) and a low-yeast diet (LYD) with half the yeast content of MYD. We noticed an inverse correlation between the number of Arc1-expressing cells in the brain and the yeast content of the diet: the number of Arc1-expressing cells increased in the SEZ and VNZ on LYD, and decreased on HYD (Figure 1B). The same changes were seen in the genetic background control *Arc^115^* line (data not shown). In agreement with the described roles of Ilp2 in response to diet, the HYD effect on Arc1 expression was similar to the effect of Ilp2 overexpression: both lowered the number of Arc1 positive cells in the brain (Figure 1A,B). In addition, when we depleted yeast from the diet and submitted the animals to a starvation regimen we observed that the levels of Arc1 in the main Arc1-producing cells were significantly higher compared to fed controls (Figure 1C). Consistent with our previous observations (Mosher et al. 2015) and a model in which yeast-responsive Ilp2 signaling regulates Arc1 expression, and not vice versa, Ilp2 levels in the brain were unaffected by *Arc1* mutation (Figure 1D).

### Arc1 mutation sensitizes larvae to dietary effects on fat storage and development

The responsiveness of Arc1 expression to diet and to Ilp2 levels are consistent with a functional role for Arc1 in maintaining energy balance in response to dietary challenges. Together with our previous work describing Arc1 as a metabolic regulator (Mosher et al. 2015), this idea led us to test Arc1 metabolic and developmental requirements under different dietary conditions, using yeast content as the challenge. We first used a sensitive buoyancy-based assay (Hazegh et al. 2017; Reis 2023; Reis et al. 2010) to measure body fat in wandering larvae reared on the HYD, MYD, or LYD. We tested a null allele, *Arc1^18^*, which is derived from an imprecise P element excision that deletes the *Arc1* coding sequence (Mattaliano et al. 2007), in comparison with *Arc1^115^* animals, which express wild-type Arc1 due to precise excision of the same P element (Mattaliano et al. 2007) and represent the genetic background control. Consistent with our previous findings (Mosher et al. 2015), *Arc1^18^*mutants accumulated more fat than *Arc1^115^* on HYD and MYD (Figure 2A). On LYD, however, the high-fat mutant phenotype disappeared and *Arc1^115^* and *Arc1^18^* showed similar levels of fat (Figure 2A). Strikingly, the fat content in both *Arc1^115^* and *Arc1^18^* animals fed the LYD appeared to increase compared to the diets with lower yeast content (Figure 2A). This paradoxical phenomenon has been observed before (Klepsatel et al. 2020; Makwisa et al. 2025) and could be explained by a delay in development caused by low dietary protein, which extends the duration of the larval feeding stage and concomitantly allows for increased consumption of this relatively carbohydrate-rich diet.

**Figure 2.**
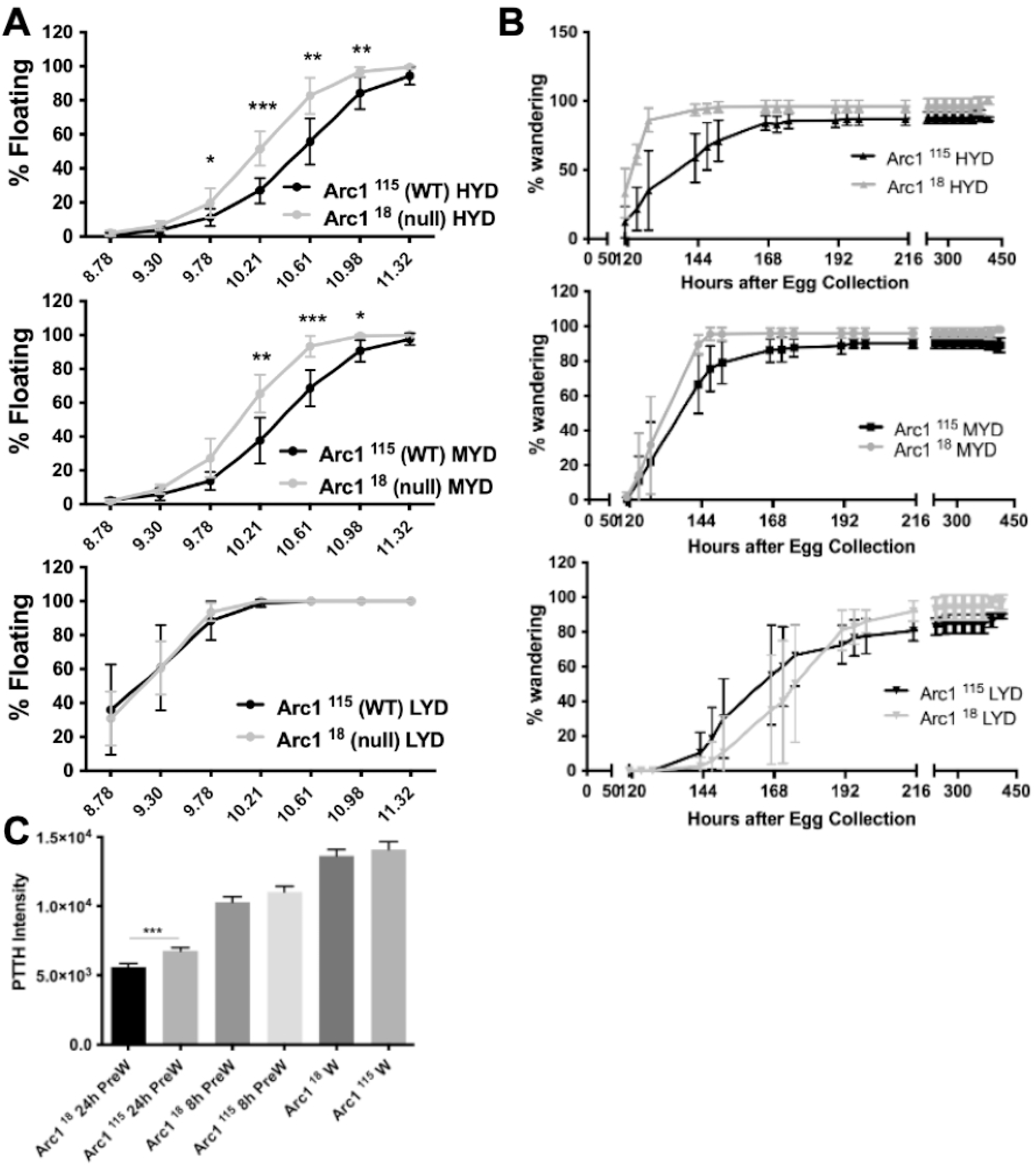
Body fat levels and development in *Arc1* mutants are highly sensitive to the nutritional content of diet. (A) Larvae of the indicated genotypes were reared on food that was rich (high-yeast diet, “HYD”), standard (medium-yeast diet, “MYD”), or poor (low-yeast diet, “LYD”) as regards the yeast content, and the fraction that started wandering was monitored at regular timepoints. Error bars, standard error of the mean. *n*=50 per genotype, 4 biological replicates. (B) Percent floating larvae at equilibrium in different density solutions for animals grown in the different diets. *n*=8 biological replicates of 50 larvae each per condition. Significant changes are observed for larvae in HYD and MYD but not LYD. P values represent results from two-tailed paired t tests. ***, 0.0001<P< 0.001; **, 0.001<P<0.01; *, 0.01<P<0.05. (C) Levels of PTTH in the PTTH-producing cells were quantified by immunohistochemistry in *Arc1^18^* mutants and control *Arc1^115^* larvae grown in MYD at 24-, 8-hrs-pre-wandering, or wandering stage. P values represent results from two-tailed paired t tests. ****P< 0.0001

Because others previously reported a developmental delay in germ-free *Arc1* mutants (Keith et al. 2021), we examined developmental timing in *Arc1* mutants under the different dietary challenges. We collected eggs from wild-type and *Arc1* mutant animals over a 4-5-hour (hr) period and ∼24 hrs later transferred first-instar larvae into vials with HYD, MYD or LYD. As expected, varying dietary yeast content altered developmental rates in both wild-type *Arc1^115^* and *Arc1^18^* animals, as measured by the number of larvae that successfully wandered, pupariated and eclosed as adults over time (Figure 2B). However, Arc1-expressing (*Arc1^115^*) animals were better able to resist changes to developmental timing when compared to the mutants. *Arc1^18^*mutants developed significantly faster than *Arc1^115^* on HYD: at 123 hr after egg deposition (AED), 61±7% of *Arc1*-mutant larvae had wandered compared to 21.5±16%, p= 0.009 for unpaired t test with Welch’s correction. There was a similar effect on MYD: at 143 hr AED, 89.5±6% of *Arc1*-mutant larvae had wandered compared to 66±17%, p= 0.049. *Arc1^18^* mutants also displayed a broader range of variability in development timing across diets. When compared to *Arc1^115^* controls, *Arc1^18^* larvae exhibited lower variability in developmental timing on HYD (*Arc1^18^* coefficient of variation (CV) was 12.4% versus *Arc1^115^* CV of 22.6%) and higher on LYD (*Arc1^18^* CV=45.7% versus *Arc1^115^* CV=39.1%). On MYD variability in developmental timing was more similar in both conditions (*Arc1^18^* CV=24.4% versus *Arc1^115^* CV=26.1%).

Based on the role of PTTH in the onset of wandering (Christensen et al. 2020) we measured PTTH levels in the PTTH-producing cells in *Arc1^115^* versus *Arc1^18^* larvae and found a significant reduction in *Arc1* mutants at the 24-hr-pre-wandering time point (Figure 2C). Reduced PTTH in these cells is consistent with a model where PTTH is released prematurely by *Arc1* mutants, which could drive earlier wandering. Together, these data show that Arc1-expressing animals exhibit a robust ability to buffer against diet-induced metabolic and developmental changes and *Arc1* mutation compromises this buffering ability.

### Diet-dependent effects of *Arc1* mutation on starvation resistance

A requirement for Arc1 in buffering organismal metabolism against dietary changes (Figure 2B) and an increase in Arc1 when nutrients are scarce (Figure 1C) suggest that Arc1 may be involved in the response to starvation. Indeed, *Arc1* mutant adults are known to be starvation resistant and Arc1 expression in IPCs restores normal starvation sensitivity (Keith et al. 2021; Mattaliano et al. 2007). To address a potential role of Arc1 during larval starvation, we measured larval survival under starvation immediately upon larval hatching. *Arc1* null mutation made no difference in survival under this condition (Figure 3). However, if we first fed larvae for 24 hr after larval hatching and then began the starvation regimen, *Arc1* null mutation prolonged survival in a manner proportional with the yeast content of the pre-starvation diet (Figure 3). These observations support a model where the metabolic defects in *Arc1* mutants (Mosher et al. 2015) drive excess storage of energy from the nutrients consumed, and the higher fat that accumulates on high-yeast diets (Figure 2A) becomes an advantage during starvation, as it provides extra energy stores from which the mutants can draw to survive when external energy sources are unavailable. Thus, while we previously showed that food intake is unchanged in *Arc1* mutants (Mosher et al. 2015), these new data demonstrate that the quality of the diet does play a major role in the metabolic phenotypes accompanying *Arc1* mutation.

**Figure 3.**
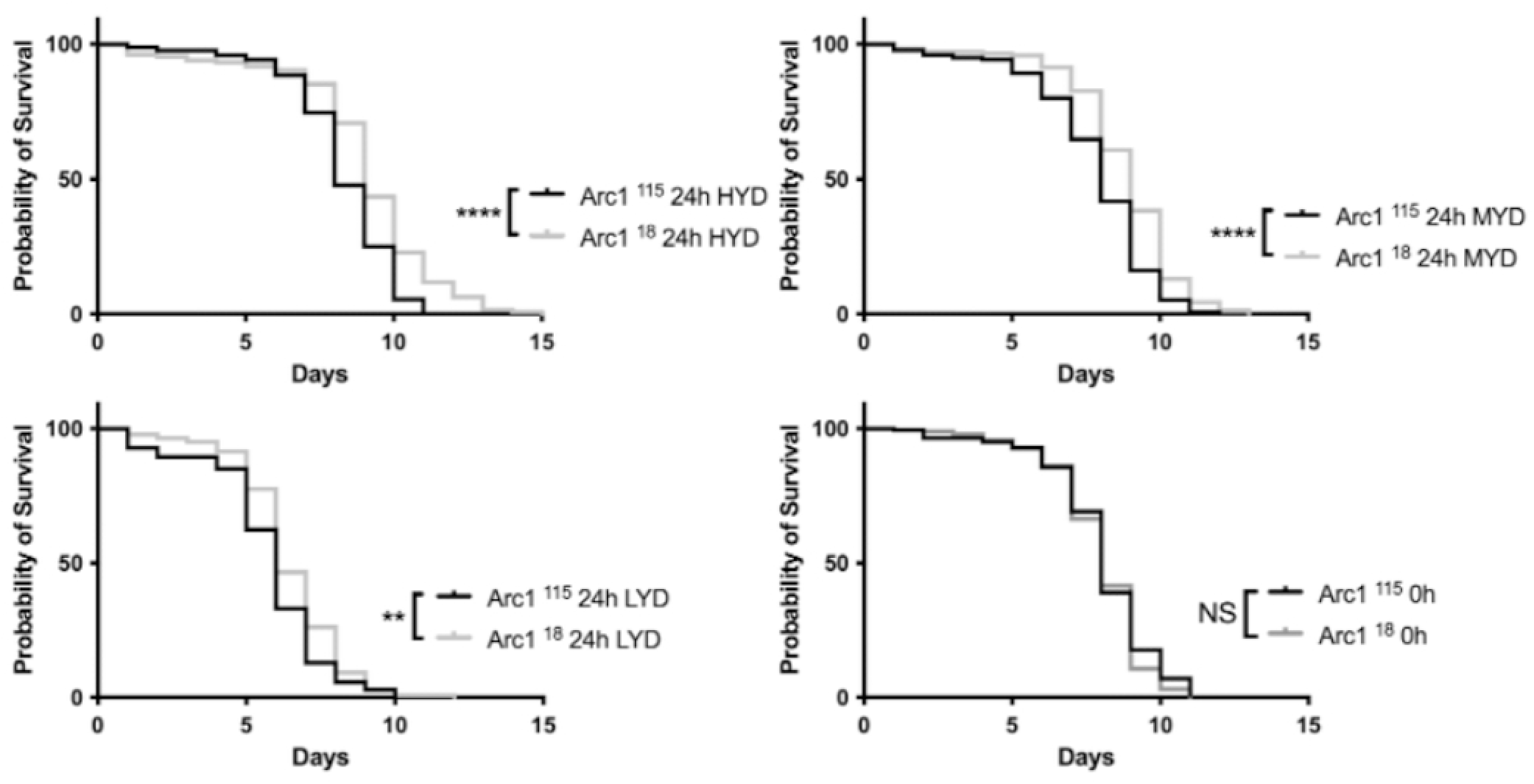
Starvation resistance in *Arc1* mutants requires prior feeding. Survival curves for larvae of the indicated genotypes fed the indicated diets for 0 or 24 hours prior to transfer to starvation medium. *n*=4 biological replicates of 45 larvae each per condition except for *Arc1^18^* in HYD, n=3 (one biological replicate was excluded due to mold contamination). Log-rank Mantel-Cox test.****, 0.00001<P< 0.0001; **, 0.001<P<0.01; NS non-significant.

### Arc1 is required for expression of genes controlling larval metabolism and development

Our data are consistent with a model in which Arc1 allows larvae to maintain rates of development close to normal despite variation in the quality of the diet. To detect for changes in the expression of metabolic or developmental genes in *Arc1* mutants that might explain the diet dependence of the fat and developmental phenotypes, we performed short-read sequencing of RNA isolated from third-instar *Arc1^18^* and *Arc1^115^* larvae fed the HYD, MYD or LYD. Bioinformatic analysis identified five clusters of genes that significantly changed expression in parallel ways (Figure 4 and Table S1).

**Figure 4.**
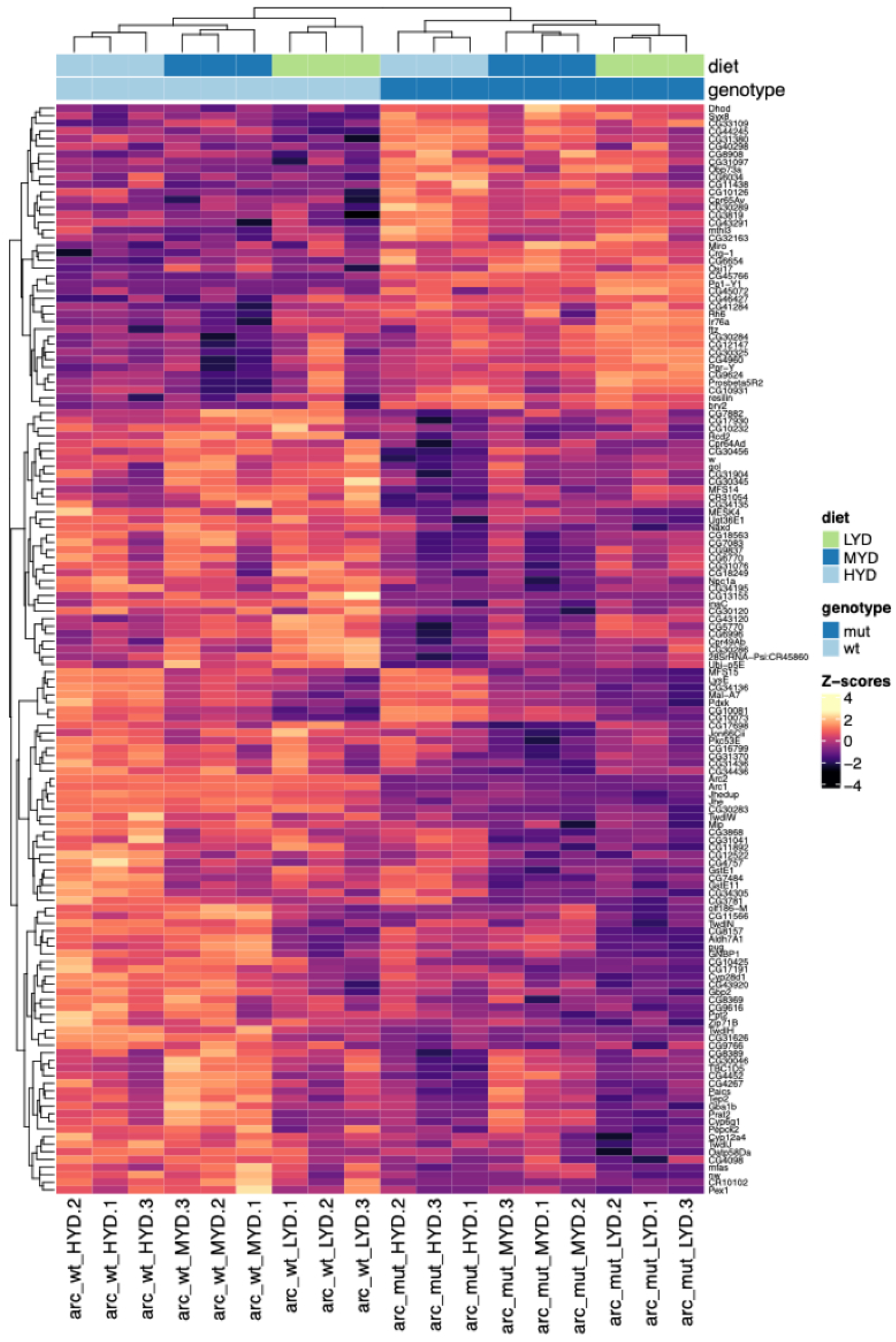
Heatmap of top differentially expressed genes in *Arc1* mutant larvae versus controls on the three different diets. Heatmap depicting 5 clusters of significantly upregulated and downregulated genes determined via Wald analysis.

We observed significant expression changes in metabolic genes, independent of diet, that provide evidence of a global metabolic defect in *Arc1* mutants. Maltase-encoding genes were a notable example (Table S1). As maltases are involved in carbohydrate breakdown, these defects in *Arc1* mutants are consistent with a defect in carbohydrate metabolism that we previously documented as manifested by high levels of glucose, trehalose and maltotriose (Mosher et al. 2015).

Consistent with the developmental differences *Arc1* mutants, we found that *Jhe* and *Jheup* (Juvenile hormone esterase duplication) both clustered tightly with *Arc1* due to their strong downregulation in *Arc1^18^* animals relative to *Arc1^115^*, regardless of diet (Figure 4). *Jhe* and *Jheup* belong to the family of carboxylesterases involved in JH catabolism and downregulation of these genes would be expected to increase JH levels (Crone et al. 2007; Hopkins et al. 2019; Liu et al. 2008). Increased JH would then counteract Ec induction of developmental progression (Santos et al. 2019). Niemann-Pick type C-1a (*Npc1a*), which encodes a cholesterol trafficking protein that contributes to sterol and ecdysone metabolism (Fluegel et al. 2006), was also downregulated in *Arc1^18^*animals on all diets (Figure 4 and Table S1). Consistent with our findings, experimental *Npc1a* downregulation increases larval ecdysone levels and alters developmental timing in a diet-dependent manner (Texada et al. 2022).

Because of the mutant-dependent, diet-specific phenotypes exhibited by *Arc1* mutants, we were also interested in genes that changed in an Arc1- and diet-dependent manner. With this goal we performed enrichment analysis using PANGEA (Pathway, Network and Gene-set Enrichment Analysis (Hu et al. 2023). Consistent with the high-fat phenotype of the mutants fed the HYD, triacylglycerol lipase activity (GO Molecular Function) was enriched (3.2-fold, p= 0.0658), reflecting downregulation of the TAG lipase genes CG18301, Lip2 and CG8093. *Arc1* mutants fed HYD showed specific enrichment (8.5-fold, p= 0.00505) of sterol biosynthesis genes (GO Biological Process), as defined by changes in CG11162 and CG1998 (both encoding 4-alpha-methylsterol monooxygenases) and *sro* (encoding a short-chain dehydrogenase/reductase). Others also found *sro* downregulation in the brains/ring glands of *Arc1* mutant larvae (Keith et al. 2021). Allatostatin A receptor 1 (*AstA-R1*) was the only other JH-responsive gene that was differently expressed in *Arc1* mutants, but in contrast to *Jhe* and *Jheup*, *AstA-R1* was only differently expressed on the HYD (Table S1). *AstA-R1* depletion from PTTH-producing cells reduces PTTH secretion and delays development (Deveci et al. 2019).

Pupariation is normally preceded by production of a glue that allows larvae to securely attach to substrates before metamorphosis (Fraenkel and Brookes 1953). The Ec-induced genes Sgs8, Sgs5, and Sgs7, which encode protein components of this glue (Beckendorf and Kafatos 1976; Da Lage et al. 2019), exhibited consistent changes in *Arc1* mutants relative to wild-type specifically in the HYD, leading to a strong enrichment in the GO Biological Function “puparial adhesion genes” (22.2-fold, p= 0.00025). Since RNA was extracted from animals collected at the same stage of development (third-instar wandering), the misexpression of puparial adhesion genes in *Arc1* mutants provides clear molecular evidence of HYD-induced alterations in developmental timing specific to *Arc1* mutants.

The gene expression differences between wild-type and *Arc1*-mutant larvae on the LYD pointed directly to Arc1 roles in synaptic plasticity. For example, *Arc1* mutants exhibited an enrichment in GO Biological processes “bulk synaptic vesicle endocytosis” (18.1-fold, p= 0.00481) due to changes in expression of *shibire* and *straightjacket* and “synaptic transmission, cholinergic” (14.8-fold, p=0.00091) due to changes in *nicotinic Acetylcholine Receptor α6*, *SLC22A family member*, and *nicotinic Acetylcholine Receptor α7*. These gene expression changes correlate strongly with the changes we observed in fat storage and rates of development in the mutants reared on diets differing in yeast content.

## DISCUSSION

Here we describe effects of *Arc1* mutation on the progression of *D. melanogaster* larval development in response to dietary changes. We further show that Arc1 expression in specific brain regions changes in response to these same dietary conditions. Our data point to a role for Ilp2 expression and secretion from IPCs in Arc1 regulation, in which yeast-dependent changes in Ilp2 (Suzawa et al. 2025) may be responsible for the changes in Arc1 brain expression we see across the diets. This effect may not be direct; other signaling mechanisms (or combinations thereof) are likely also at play. What mediates diet regulation of Arc1 function in the brain is an exciting area of future research. We note that whereas we saw increases in the number of specific larval brain cells expressing Arc1 when we decreased the nutritional content of the diet (Figure 1B,C), the microbiota study reported a starvation-induced decrease in overall Arc1 levels in adult heads (Keith et al. 2021). Beyond possible differences between larvae and adults, we focused on specific brain regions and did not measure overall brain Arc1 levels.

What is clear is that in both cases brain levels of Arc1 respond to diet. Also unlike our results (Figure 2D), Keith et al. found changes in Ilp2 protein levels in IPCs of *Arc1*-mutant larvae compared to wild-type. As Ilp2 responds to yeast content of the diet (Suzawa et al. 2025) and the yeast content in our diets is different from theirs, differences in diet composition may explain this apparent discrepancy.

The study centered around roles of Arc1 in the microbiota also found diet-dependent changes in development in *Arc1* mutants, but these changes were exclusively delays and were only observed in animals lacking gut bacteria (Keith et al. 2021). The effect of gut bacteria was ultimately attributed to effects of a single bacterial genus, *Acetobacter*, on the food. Thus, by altering what nutrients are available as ingested food is processed in the gut, the microbiota effectively modifies the diet. We saw on a high-yeast diet that *Arc1* mutants developed faster than the genetic background control. The microbiota study found that, on most diets, gnotobiotic larvae (i.e., with gut microbiota present) developed at similar rates regardless of *Arc1* mutation. Only on two specific diets (5% yeast, 3% glucose and 3% yeast, 5% glucose) were developmental differences seen in gnotobiotic *Arc1* mutants, and development was slower, not faster (Keith et al. 2021). However, those diets are qualitatively and quantitatively different than ours, hence direct comparison is not possible. Nonetheless, we find it noteworthy that among all the various conditions in that study the animals showing the fastest (in 5% yeast 5% glucose) and those showing the slowest rates of development (in 3% yeast 10% glucose) were Arc1 mutants (Keith et al. 2021). Additionally, genetic background of the different Arc1 mutants also likely contributes to differences with the microbiota study: in the Oregon R background, a pseudogene located between *Arc1* and *Arc2* correlates with a drastic reduction in Arc1 expression (Ashley et al. 2018). In “wild type” animals of the Oregon R background, synaptic plasticity (as assessed by the number of synaptic boutons at the neuromuscular junction) is reduced relative to Canton-S, which lacks the pseudogene (Ashley et al. 2018).

We have previously reported with another gene (Gillette et al. 2020) gene-diet interactions that “rescued” mutants from developmental delays observed on the standard diet. In that situation, the positive effects of the altered diets were more pronounced in the mutants than the wild type, but wild-type animals always developed faster than the mutants; diets could at best reduce the magnitude of the genotypic effect on the phenotype (Gillette et al. 2020). Here, we find that without Arc1 the effects of diet can be more profound: on some diets, Arc1 mutants developed faster than did the wild type, inverting the direction of the genotypic effect.

Arc1 levels in the brain increased in animals fed the LYD (Figure 1B), consistent with a higher functional requirement for Arc1 in the brain in this dietary condition. Indeed, on this diet we saw a clear enrichment for genes involved in brain- and neuron-related processes among those that were significantly different between wild-type and *Arc1*-mutant animals (Table S1). *Arc1*-mutant larvae fed LYD developed more slowly than wild type (Figure 2B), presumably due to changes in the expression of genes involved in JH synthesis. For example, diet-independent defects in *Arc1* mutants in the production of Jhe and Jheup –– two esterases involved in JH degradation –– may result in an altered regulatory feedback loop between JH and Ec, leading to smaller, broader peaks of Ec that render larvae more vulnerable to dietary influences on developmental timing.

By contrast, on HYD, where *Arc1* mutants develop faster than wild type, we saw differences between wild-type and *Arc1*-mutant larvae in the expression of genes related to sterol synthesis and lipid metabolism (Table S1). Under such rich conditions, where sterol metabolism is differentially regulated and nutrients are in excess, *Arc1*-mutant larvae can likely compensate and overcome the inhibitory effects of JH on Ec, resulting in faster development. In these animals, there is less Arc1 in the brain (Figure 1B), consistent with a model in which Arc1 functions in energy metabolism outside the brain dominate on a nutrient-rich diet. *Drosophila* Arc1 and human ARC appear to have evolved from independent “domestication” events of the Gag domain of a retrotransposon (Ashley et al. 2018; Pastuzyn et al. 2018; Shepherd 2018). Arc1 and human ARC proteins retain the ability to oligomerize into capsid-like particles and to package RNA, including the dArc1 mRNA, into those particles, which are released from neurons in extracellular vesicles. In flies, these vesicles are received by muscle cells across the neuromuscular junction and mutations that block this process perturb synaptic plasticity (Ashley et al. 2018). Arc1-containing capsids could mediate communication with energy storage tissues via a similar mechanism. Overall, our data support a model that centers Arc1 as a key coupling factor that responds to levels of nutrient availability via Ilp2, ultimately regulating in expression of metabolic and developmental genes and resulting in a regulatory mechanism integrating developmental timing with dietary and metabolic needs.

## Materials and methods

### Fly Strains and husbandry

w^1118^ (stock number 3605) was obtained from the Bloomington stock center. *Arc1^115^*, *Arc1^18^*, were a generous gift from Leslie Griffith, and were backcrossed to the w^1118^ stock. *Arc1^18^* is null mutant resulting from an imprecise excision allele of a P-element insertion at the Arc1 locus, which deletes the Arc1 coding sequence without affecting surrounding genes (Mattaliano et al., 2007). *Arc1^115^* is a precise excision allele that restores the full Arc1 coding sequence and is used as the genetic background controls for *Arc1^18^* (Mattaliano et al., 2007). Larvae and flies were reared as previously described (Reis et al., 2010). Unless otherwise specified, all animals were reared at 25°C, 60% humidity and fed Bloomington media with malt extract (Reis et al., 2010). Eggs were collected on grape agar plates for 4-5 hrs and ∼24 hrs later 50 first instar-larvae per biological replicate were transferred to the wide-media vials (Genesee cat# 32-117) containing the experimental diets.

### Diets

Experimental media was made fresh each week and used for no longer than that week. All diets contained, per L: 9.2 g soy flour (Soy Flour Defatted, ADM Specialty Ingredients, cat. # 63-100), 65 g yellow cornmeal (Quaker Yellow Corn meal enriched degerminated), 42.4 g light malt extract (Briess Gold Light Dry Malt Extract Cat. # 574*8)*, 5.3 g agar (Apex Chemicals cat. # 66-104), 70 mL light corn syrup (Karo Light Corn Syrup), 4.4 mL propionic acid (J.T.Baker, CAS 79-09-4), and 8.4 mL Tegosept (Apex Bioresearch Products #20-258, 380 g in 1L 100% ethanol). Yeast (Fleischmann’s Instant Dry Yeast #2139) content per L: 70g for HYD, or 35g for MYD, or 7g for LYD.

### Starvation regimen

Animals were fed in until ∼84 hrs after egg deposition, then transferred to agar/water media (5.3 g agar per liter, same as food) for starvation, or to a new vial of regular food as control, for 12hrs, then dissected and used for immunohistochemistry. n= 50 brains total, 3 biological replicates.

### Density Assay

Density assays were performed as previously described (Hazegh and Reis 2016; Reis et al. 2010). All experimental conditions and genotypes were analyzed with 8 independent biological samples, of 50 larvae each. Two-tailed paired t test was used to calculate statistical significance with GraphPad Prism software test.

### Developmental Timing

Number of animals at each developmental stage (wandering larvae, pupated or eclosed adults) was recorded every day at 10:00, 14:00 and 18:00. n=50, 4 independent biological replicates.

### Microscopy and cell counts

Wandering 3rd instar larvae were dissected by inversion and fixed in a microcentrifuge tube with 4% paraformaldehyde (Electron Microscopy Sciences, Cat # 15710) in PBS for 30min at RT in a nutating mixer. Carcasses were then washed 3 times with 0.1% PBTriton and immunostaining performed as described (Reis et al., 2010). Arc1 antibody (Mattaliano et al., 2007) was used at a 1:500 dilution and was a generous gift from Leslie Griffith. Notably, this Arc1 antibody has been tested extensively for specificity, and immunostaining signals are absent in null animals ((Mattaliano et al., 2007) and our unpublished results). Primary antibodies were incubated 4°C. Ilp2 antibody was used at 1:800 (Geminard et al. 2009) and PTTH antibody was used at 1:500 (Yamanaka et al. 2013b); both were generous gifts from Pierre Leopold. Secondary antibodies were incubated for 2 hrs at RT (anti-rabbit Alexafluor 488 (Invitrogen cat# A11034) anti-rat Alexa Fluor 568 (Invitrogen cat# A11077) and anti-guinea pig Alexa Fluor 568 (Invitrogen cat# A11075)) were used at 1:5000 dilution. After final washes carcasses were dissected and isolated brains were mounted on slides with Slowfade Gold Antifade Mountant (Invitrogen, cat# S36936) and stored at −20° until imaging. Images were collected using a Leica TCS SP5 II confocal microscope. For cell counts (Figure 1B) data were blinded prior to collection and analysis. The number of cells positive for Arc1 antibody staining was determined by manual counting. Fluorescence intensity was calculated using FIJI/ImageJ (Schindelin et al. 2012) by subtracting the integrated density of an area with no fluorescence from the integrated density of a region of the same area. Multiple t test comparison was used to calculate statistical significance, using Prism 6 software. Stainings were performed from 3 independent biological replicates and at least 50 brains analyzed per sample.

### Starvation resistance

Crosses of the appropriate genotype were set up for egg collection without yeast paste. Upon larval hatching, animals were transferred to an agar media vial (starvation) or into a vial with the designated diet. 24 hrs later, all animals (starved and fed) were transferred to vials with agar-only starvation medium by adding a solution of 20% sucrose to float the larvae to the surface. Number of live and dead animals were counted daily at approximately the same time in the afternoon. n=45 larvae, 4 biological replicates each. One of the biological replicates *for Arc1^18^*in HYD was excluded from analysis due to mold and bacteria contamination.

### RNA sequencing

*Sample preparation:* Total RNA was extracted from 50 wandering third-instar *Arc1^115^ or Arc1^18^* using Trizol reagent (Life Technologies cat# 15596026) following the manufacturer’s instructions. RNA sequencing and library prep was performed at the University of Colorado Anschutz medical campus Genomics Core.

### Library Preparation

Libraries were prepared according to the manufacturer’s protocol using the Universal Plus mRNA-Seq library preparation kit with NuQant (TECAN Cat # 0520-24).

### Data acquisition and processing

Libraries were sequenced with an Illumina NovaSEQ 6000 system. Transcriptome analysis was performed using pseudo-alignment with Salmon using the *D. melanogaster* transcriptome. DESeq2 was used for differential expression analysis. PANGEA (Pathway, Network and Gene-set Enrichment Analysis (Hu et al. 2023) was used for the enrichment analysis of differential expressed classes of genes in mutant versus control across the different diets.

## Data availability statement

The data underlying this article are available in the article and in its online Supplementary material. Raw gene expression data are available at GEO.

## Acknowledgements

We thank FlyBase (Aguila et al. 2007; Jenkins et al. 2022; Ozturk-Colak et al. 2024) for providing a comprehensive database with data, tools and resources critical for the design and execution of this project, and the Bloomington Drosophila Stock Center, supported by grant NIH P40OD018537, for generously maintaining and providing fly stocks. We thank Leslie Griffith for generously sharing fly stocks and the Arc1 antibody, and Pierre Leopold for sharing the ILP2 and PTTH antibodies. Michael McMurray helped editing the manuscript.

## Author contributions

W.Z. data collection and analysis (Figures 1-3); L.S. data collection and analysis (Figure 2) and data collection (Figure 4); K. R. RNA sequencing data analysis; T.R. conceptualization, methodology, data collection and analysis (Figure 4), formal analysis, validation, funding acquisition, supervision, writing.

Table S1. RNA-sequencing analysis of *Arc1* mutant larvae versus controls on the three different diets.

